# SVelfie: A method for discovering cancer drivers based on enrichment of likely functional structural variants

**DOI:** 10.1101/2025.03.09.642284

**Authors:** Xavi Loinaz, Johnathan Dagan, Jean-Baptiste Alberge, Antonia Kowalewski, Mendy Miller, Julian M. Hess, Esther Rheinbay, Chip Stewart, Gad Getz

## Abstract

Structural variants (SVs) can drive tumorigenesis, yet discovering SV cancer drivers remains challenging^1^. Here, we present SVelfie (**<u>S</u>**tructural **<u>V</u>**ariants **<u>e</u>**nriched with **l**ikely **<u>f</u>**unctional/**i**mpactful **<u>e</u>**vents), a statistical method to infer driver genes from SVs detected across a cohort of cancer genomes, based on enrichment of likely functional events. When testing SVelfie on lymphoma samples from the Pan-Cancer Analysis of Whole Genomes (PCAWG)^2^, it corroborates known tumor suppressor genes and also yields novel driver candidates.

## Main

Cancer genomes often contain 100s to 100,000s of somatic genetic alterations^3^, including point mutations (single-nucleotide variants [SNVs] and short insertions/deletions [indels]), copy number variants (CNVs), and structural variants (SVs). Only a small number of these alterations drive tumorigenesis^4^, making the inference of such driver events and the corresponding genes a challenging task. Due in part to the historically greater availability of whole-exome sequencing (WES) relative to whole-genome sequencing (WGS) data, many robust methods have been developed for inferring driver genes from somatic point mutations/indels and CNVs, both of which can be inferred from WES data from tumors and their patient-matched normal DNA^5–9^. To detect SVs comprehensively, however, WGS data is necessary because most SV breakpoints are located in regions outside of exons^10^. Historically, WGS has been prohibitive due to high cost, limiting both the number of discovered SV drivers and the development of robust methods to find them. This highlights the current need to identify cancer-driving SVs, especially due to the functional effect and driving potential of SVs on cancer initiation and/or progression^11–13^, which can often exceed that of point mutations or CNVs^14–16^, the latter of which are typically also accompanied by SVs at their boundaries^17^.

With the ongoing decrease in cost of sequencing, and the resulting increased availability of WGS cancer data sets, we now have greater statistical power than ever to discover cancer drivers based on SVs. This opportunity to detect SV drivers will only further widen over time, creating opportunities to identify functionally relevant rearrangements that can ultimately improve cancer diagnosis, treatment, and the identification of novel drug targets. Thus, there is a growing need for developing robust, novel and complementary methods for discovering SV drivers.

Several methods are currently available to discover SV drivers, such as CSVDriver^18^, fishHook (for 1D breakpoint analysis^12,19^), and SVSig (for 2D adjacency analysis^12,20^). These methods identify candidate driver genes, genomic regions, or adjacencies that are more frequently hit by SV breakpoints than expected from a model of background “passenger” structural variants. Each method uses a different approach and distribution to estimate the expected frequency (and its uncertainty) of background breakpoints. Modeling such background frequencies (or densities) can be difficult since SVs are relatively rare events (in PCAWG there were on average two orders of magnitude fewer SVs than point mutations^2^), and the genomic covariates that affect them are not well understood^21^. Moreover, when trying to assess the frequency of SVs connecting two specific loci in the genome (“adjacencies”), the search space becomes even larger, quadratically scaling with the size of the genome, further increasing the problem of sparsity. To overcome this sparsity issue, one can instead compare, for every gene, the observed fraction of SV events associated with the gene that have specific functional effects to the fraction expected by chance. This approach is similar to methods for discovery of point mutation drivers which search for genes with an unexpectedly high ratio of non-synonymous (deleterious) to synonymous (neutral) mutations (dN/dS)^5^, regardless of the total number of mutations in the gene. Since this approach does not depend on the total number of SVs affecting a gene (or region) and is thus complementary to the aforementioned background density-based methods, it can be used either on its own or in conjunction with the density-based methods.

We present such a functional enrichment-based approach, which we call **SVelfie (<u>S</u>**tructural **<u>V</u>**ariants **<u>e</u>**nriched with **l**ikely **<u>f</u>**unctional/**i**mpactful **<u>e</u>**vents; pronounced “*SVEL-fee*”). SVelfie finds genes that are enriched with SVs that presumably cause likely loss-of-function (LLoF), analogous to the dN/dS approach for point mutations.

## Results

### Overview of the SVelfie framework

SVelfie is a statistical method that infers SV drivers by identifying enrichment of LLoF events for candidate genes, using an orthogonal approach complementary to background density-based SV driver discovery methods. Specifically, it enables discovery of putative tumor suppressor genes which have a higher-than-expected fraction of SVs that likely cause loss-of-function compared to neutral events. SVelfie’s two most crucial steps are to (a) estimate the expected fraction of LLoF SV events for each tested gene, and (b) find genes with statistically significant enrichment of LLoF based on that expected fraction (corresponding to steps [ii] and [iii] in **Figure 1A**). Both of these steps rely on our designation of LLoF and unlikely loss-of-function (ULoF) events (**Figure 1B**; **Methods**), as well as how we associate SV breakpoints to gene transcripts and annotate them. Breakpoints are annotated using a modified version of the annotatation step of dRanger^22^, a well-established SV detection method (**Methods**).

**Figure 1:**
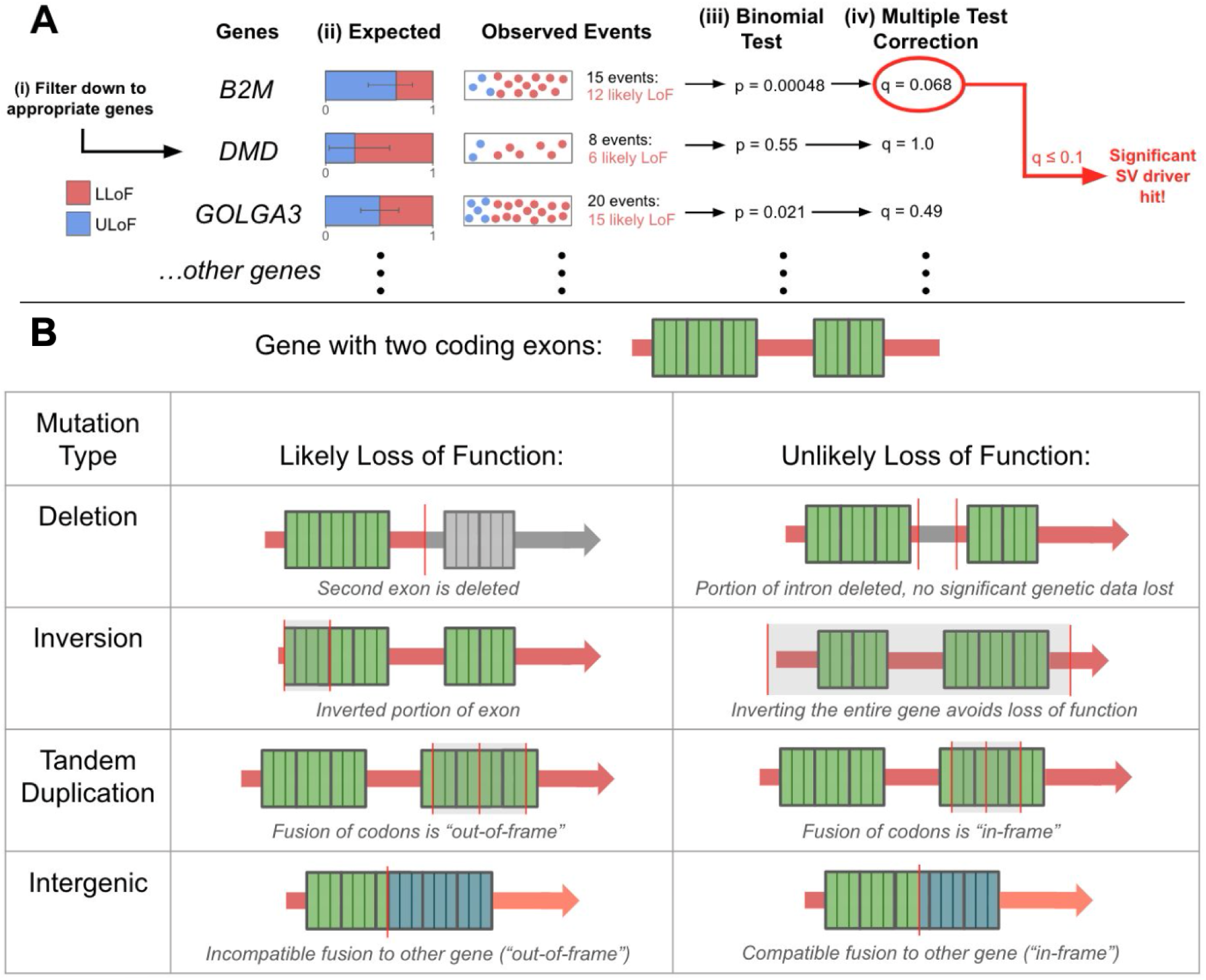
Overview of the framework and conventions for SVelfie. **A. SVelfie analysis steps:** (i) Filter down genes to those with potential for being significantly enriched in LLoF; (ii) Calculate the expected fraction of likely loss-of-function (LLoF) SVs for each gene; (iii) Calculate the significance of the observed number of LLoF events (p-value); (iv) Correct for multiple hypothesis testing using the Benjamini-Hochberg procedure (q-value). Genes below a specified q-value threshold are considered candidate SV drivers (**Methods**). **B. Graphic representing examples of our approach to determine LLoF events and events that are unlikely loss-of-function (ULoF).** In the example gene, there are three codons in the first exon and two codons in the second exon, with each codon having bolder borders than individual nucleotides. Full descriptions of the mapping of SV configurations around a gene to LLoF or ULoF are in **Methods**.

### SVelfie identifies known and novel candidate LLoF SV drivers in lymphoma

In order to test SVelfie’s ability to infer SV drivers, we applied SVelfie to SV calls available from the Pan-Cancer Analysis of Whole Genome (PCAWG) project^2^ of the International Cancer Genome Consortium (ICGC) and The Cancer Genome Atlas (TCGA). We used the 107 available lymphoma cases (**Methods**) since they have known SV drivers that can serve as positive controls.

SVelfie identified 10 candidate tumor suppressor genes (**Figure 1**; **Table 1**). Overall, 5 out of 10 of these candidate drivers (*CDKN2A, MTAP, CREBBP, CDKN2B, FBXO11*) are well-known as tumor suppressor genes in lymphoma. *CDKN2A* (SVelfie q=9.4x10^-6^) is one of cancer’s most studied tumor suppressor genes^23^; it is also reported as having recurrent deletions in follicular lymphoma^24^, diffuse large B-cell lymphoma (DLBCL)^25^, and peripheral T-cell lymphoma not otherwise specified (PTCL-NOS)^26^. *CDKN2B* and *MTAP* are adjacent to *CDKN2A,* and this region is often lost by focal copy number alterations (hence the SVs in this region). *CDKN2B* is a tumor suppressor gene in its own right^27^ and often acts in tandem with *CDKN2A* to regulate the cell cycle^28^. *MTAP* loss may be a bystander effect of deleting *CDKN2A*/*B*, but it leads to accumulation of methylthioadenosine (MTA), which then results in a dependency on PRMT5 and was recently targeted by PRMT5 inhibitors^29^. *CREBBP* is a well-established tumor suppressor gene in lymphomas, frequently inactivated by mutations or deletions in follicular lymphoma and diffuse large B-cell lymphoma^30,31^. As a histone acetyltransferase, *CREBBP* plays a critical role in regulating transcription and maintaining normal immune cell differentiation, with its loss contributing to impaired tumor suppressive pathways and increased lymphomagenesis^32^. Finally, loss of *FBXO11* is known to lead to stabilization and accumulation of BCL6^33^, a well-known oncogenic driver for B-cell lymphoma^34,35^. Reassuringly, these 5 driver genes are also listed as tumor suppressor genes in OncoKB^36^.

**Table 1:** Summary of SVelfie’s candidate hits (q≤0.25) on the PCAWG lymphoma cohort.

| Gene | Number of tumors | Number of tumors with at least one LLoF event | Number of events | Number of intragenic events | Number of intergenic events | Number of LLoF events | Expected frequency of LLoF | p-value | q-value |
| --- | --- | --- | --- | --- | --- | --- | --- | --- | --- |
| *KLF12* | 6 | 2 | 30 | 2 | 28 | 22 | 0.216 | 2.017E-09 | 8.894E-07 |
| *CDKN2A* | 22 | 16 | 49 | 24 | 25 | 21 | 0.118 | 4.272E-08 | 9.420E-06 |
| *CD70* | 12 | 6 | 15 | 10 | 5 | 7 | 0.113 | 6.600E-04 | 9.702E-02 |
| *MTAP* | 23 | 13 | 52 | 25 | 27 | 18 | 0.163 | 1.018E-03 | 1.038E-01 |
| *CREBBP* | 8 | 5 | 9 | 7 | 2 | 6 | 0.168 | 1.177E-03 | 1.038E-01 |
| *CDKN2B* | 22 | 12 | 49 | 24 | 25 | 13 | 0.111 | 2.053E-03 | 1.509E-01 |
| *HVCN1* | 5 | 4 | 6 | 6 | 0 | 4 | 0.131 | 3.499E-03 | 2.185E-01 |
| *TOX* | 7 | 6 | 12 | 5 | 7 | 7 | 0.201 | 3.963E-03 | 2.185E-01 |
| *RIMS1* | 5 | 3 | 14 | 1 | 13 | 9 | 0.276 | 4.512E-03 | 2.211E-01 |
| *FBXO11* | 7 | 5 | 9 | 6 | 3 | 5 | 0.147 | 5.072E-03 | 2.237E-01 |

Among the remaining 5 hits (*KLF12, CD70, HVCN1, TOX, RIMS1*), *TOX* and *CD70* have been previously described as recurrently altered via inactivating lesions, as well as nonsense and frameshift mutations in DLBCL. *TOX* has been found to be a significant biomarker for various lymphomas^37–39^, and *CD70*-inactivating lesions have been suggested to lead to immune escape in DLBCL^40^. Lower *CD70* expression has also been associated with poor overall survival in DLBCL^41^. Another one of the 5 genes, *HVCN1*, is consistent with being a tumor suppressor, as its expression was correlated with nonproliferative status for DLBCL cases^42^. *HVCN1* mutations were also found to be associated with, and clinically relevant to, follicular lymphoma^43^, and although their functional effect was not readily apparent, such mutations were predominantly frameshift, nonsense, or splice-donor mutations, which usually lead to gene function loss. Together with the LLoF evidence from SVelfie, these data suggest that *HVCN1* is a tumor suppressor in DLBCL.

The two remaining candidate drivers, *KLF12* and *RIMS1*, have hits in only a few tumors with LLoF SV events (2 and 3, respectively), with a single tumor contributing a large number of LLoF events. We would need additional orthogonal evidence for these genes in order to support their roles as tumor suppressors.

### SVelfie is well-calibrated and robust

To validate that the statistical framework of SVelfie is well-calibrated, we leveraged the fact that most genes are not SV drivers and compared the observed distribution of p-values to the expected uniform distribution of p-values for “passenger” genes (i.e., those that follow the null hypothesis) by inspecting the quantile-quantile (Q–Q) plot. For visualization purposes, we used a p-value drawing method that takes into account that the SVelfie p-values are discrete, as they are based on integer numbers of SVs (**Methods**). The Q–Q plots (**Figure 2B**) suggested that the p-values are mostly well-calibrated, with some p-value inflation likely due to overdispersion not captured by the current model.

**Figure 2:**
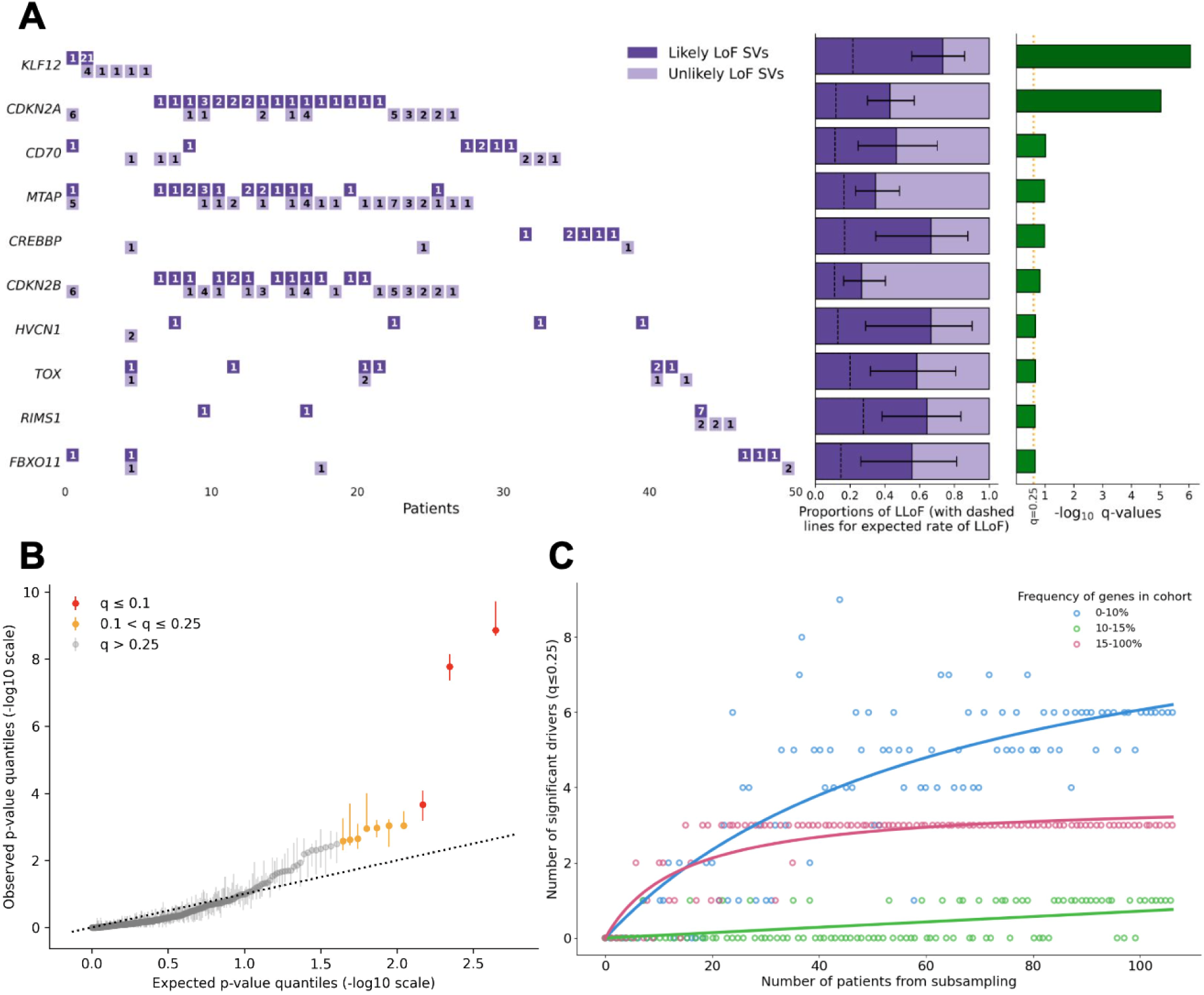
SVelfie successfully identifies known and new candidate SV drivers from the PCAWG lymphoma cohort. **A. Overview of SVelfie’s SV driver hits**. Rows represent genes with BH-FDR q≤0.25 and columns represent patients. For each gene and patient, the number of likely LLoF and ULoF SV events are shown. In the middle, the observed (with 95% confidence interval) and expected (dashed line) fraction of LLoF events across the cohort are shown (**Methods**). On the right, the significance level (q-value) for each gene is shown with a dotted line at q=0.25. **B. Q–Q plot comparing SVelfie’s distribution of p-values to a uniform distribution.** Vertical lines indicate the range of possible p-values for each gene, due to the discrete nature of the test (**Methods**). **C. Down-sampling saturation analysis on candidate driver genes yielded by SVelfie.** Genes are stratified by their frequencies across the cohort, to which Michaelis-Menten curves are fitted.

Next, we performed a down-sampling saturation analysis to investigate the effect of the number of patients on the list of candidate cancer genes. This type of analysis can indicate whether increasing the cohort size will likely yield new candidate drivers, and follows the analysis in Lawrence et al.^9^ for discovering point mutation drivers using MutSig2CV. With the number of samples analyzed here, saturation was achieved for discovering common drivers that are associated with SVs in ≥10% of cases (**Figure 2C**). However, this dataset is underpowered to discover drivers associated with SVs in <10% of cases. An extrapolation of the down-sampling analysis suggests that many more SV drivers in this frequency range are yet to be discovered.

Since intrachromosomal CNVs are associated with SVs at their boundaries, we next wanted to test whether the candidate SV drivers we detected could also be found by searching for recurrent CNVs. We therefore compared SVelfie’s significant gene list with GISTIC2.0 peak regions^7^ previously found for lymphoma-associated cohorts in PCAWG. Of the 10 SVelfie hits, only two overlap GISTIC wide-peak regions (*CD70* and *CDKN2A*) for such cohorts (specifically the “Hematopoietic_tumors”, “Lymph-BNHL”, and “Lymph_tumors” cohorts [**Table 2**]) and two are immediately adjacent (*CKDN2B* and *MTAP*). This suggests that SVelfie can capture independent evidence for SV driver detection beyond recurrent copy-number alterations.

**Table 2:** Auxiliary information for validation of SVelfie’s hits. TSG stands for tumor suppressor gene. CGC stands for Cancer Gene Census^44^ ‒ a curated catalog of cancer genes. Since 104 of the 107 tumors had RNA-seq data, the numbers in the last column may slightly differ from the numbers in **Table 1**.

| Gene | Is TSG in CGC? | Is TSG in OncoKB? | Related PCAWG cohorts with corresponding GISTIC deletion peak | Gene cytoband | RNA Wilcoxon depletion test p-value | RNA Wilcoxon depletion test q-value | Tumors associated to gene in RNA Wilcoxon depletion test |
| --- | --- | --- | --- | --- | --- | --- | --- |
| *KLF12* | no | no | None | 13q22.1 | 7.348E-01 | 7.348E-01 | 6 |
| *CDKN2A* | yes | yes | 'Hematopoietic_tumors', 'Lymph-BNHL', 'Lymph_tumors' | 9p21.3 | 6.721E-09 | 6.721E-08 | 21 |
| *CD70* | no | no | 'Hematopoietic_tumors', 'Lymph-BNHL', 'Lymph_tumors' | 19p13.3 | 5.177E-02 | 1.726E-01 | 12 |
| *MTAP* | no | yes | None | 9p21.3 | 1.147E-01 | 2.367E-01 | 22 |
| *CREBBP* | yes | yes | None | 16p13.3 | 1.183E-01 | 2.367E-01 | 8 |
| *CDKN2B* | no | yes | None | 9p21.3 | 2.718E-01 | 4.530E-01 | 21 |
| *HVCN1* | no | no | None | 12q24.11 | 6.048E-01 | 6.720E-01 | 5 |
| *TOX* | no | no | None | 8q12.1 | 5.797E-01 | 6.720E-01 | 7 |
| *RIMS1* | no | no | None | 6q13 | 5.604E-01 | 6.720E-01 | 5 |
| *FBXO11* | yes | yes | None | 2p16.3 | 2.310E-02 | 1.155E-01 | 6 |

Since SVelfie identifies tumor suppressor genes with LLoF events, we further examined whether we could observe a decrease in RNA expression of the putative driver genes in samples that contained associated SV events compared to samples that did not. Indeed, 5 of the 10 SVelfie hits (*CDKN2A*, *FBXO11*, *CD70*, *MTAP*, *CREBBP*) had significantly lower expression in cases with SVs associated with these genes compared to cases without (Wilcoxon Rank-Sum test followed by Benjamini-Hochberg for multiple hypothesis testing among candidate hits; q≤0.25 [**Table 2; Methods**]). This supports that SVelfie can identify driver genes that are indeed hindered in their expression, and thus, also likely, in their function.

## Discussion

We here present SVelfie, a statistical method for discovery of SV drivers that, to the best of our knowledge, takes a novel approach by finding significant enrichments of SV types and breakpoint configurations that likely lead to functional outcomes. Because such an approach is orthogonal to existing SV driver discovery approaches that are based on comparing the observed frequency of SV breakpoints (or adjacencies) to an estimated background density, SVelfie can complement these other approaches. In particular, since SVelfie’s statistical analysis assumes that the total number of observed SV events is fixed, the significance levels (p-values) of SVelfie and a density-based method are statistically independent and, therefore, could potentially be combined to yield a more powerful driver discovery test (e.g., using Fisher’s p-value combination method).

For this initial version of SVelfie, we focused on enrichment of LLoF SV events, and provided evidence that SVelfie not only recapitulates well-known drivers for lymphomas (using data from PCAWG), but also infers lesser-known candidate drivers, which warrant further investigation. The SVelfie statistical test is largely well-calibrated, with the p-values of the majority of genes following the uniform distribution that is expected from “passenger” genes (i.e., those following the null hypotheses). Moreover, SVelfie is robust, as shown by the saturation analysis; in the lymphoma cohort of 107 patients, it can robustly detect drivers when run on subsets of the data. SVelfie also finds SV drivers that are not significant by recurrent copy number alterations. This underscores the utility of methods based on SVs for finding novel drivers (e.g., ones affected by genomic inversions). Finally, we showed that RNA-seq evidence can corroborate SVelfie’s results.

SVelfie has several limitations that could impact its accuracy in identifying SV driver genes. First, since the statistical power of SVelfie increases with the number of observed SV events in a gene, which grows with cohort size, SVelfie is suitable for the analysis of larger cohorts, in which many genes (both drivers and passengers) are associated with multiple SV events. Due to dropping sequencing costs and resultantly more tumors being sequenced, we expect the sizes of the cohorts with SV data to dramatically increase in the near future. In comparison, the statistical power of background density-based driver discovery methods also increases with sample size, but such methods could potentially detect drivers, even if they have few SVs, when the estimated background density is low. Second, SVelfie’s statistical test is calculated based on the fraction of LLoF SVs rather than the fraction of tumors with LLoF events. This can lead to over-significance of a gene in the case that a single complex SV event is recognized as multiple independent LLoF events affecting the same gene. We plan to address this in future versions. Third, adjacent or overlapping genes may be declared as independent candidate drivers since the same SV event can be associated with multiple genes. Therefore, we recommend identifying neighboring genes in the list of candidate drivers (e.g. *CDKN2A*/*B* and *MTAP* in our DLBCL analysis) and using orthogonal data (e.g. RNA-seq or experimental data) to provide further evidence to identify the true driver(s). Lastly, SVelfie relies on conventions for what specific LLoF events look like. It is possible for SV events to be incorrectly labeled as LLoF when, in reality, a gene may not lose its function, and could even increase its expression or gain novel functionality. One example is the deletion of exon 19 in *EGFR*, which would be considered LLoF by SVelfie’s convention. However, this deletion is well-known to lead to a constitutively active protein, causing cell proliferation^45^.

Overall, SVelfie demonstrates a novel, robust approach for SV driver inference that can complement existing SV driver inference methods, recapitulating known SV drivers and yielding novel candidates. We have also applied SVelfie to a separate cohort of >1,000 multiple myeloma cases, finding 14 candidate drivers (Alberge,* Dutta,* Poletti,* et al., in print). Inference and discovery of SV drivers yields insights into cancer genes and pathways, and open up new avenues for improved cancer therapies and future treatment strategies.

## Methods

### Defining types of SV events

For each gene in the genome, SVelfie looks at all SV events that have at least one breakpoint contained within the gene or within 1 Mb from either edge of the gene (where edges are defined as the most extreme 5’ and 3’ bases of any of the gene’s transcripts used for annotation and a 3 kb imputed promoter region). As we search for SVs that can directly affect the gene, we chose a window size of 1 Mb following the expected range for cis-acting pQTLs^46^. We denote intragenic SVs as those wherein both breakpoints are local, within the 1 Mb window around the gene, and intergenic SVs as those where only one breakpoint is within the 1 Mb window. We also have three subclassifications for intragenic SV events –– deletions, inversions, and tandem duplications –– that are based on the orientation of the discordant reads supporting the breakpoint (**Table S1**).

**Table S1:** Types of SV events and their definitions. The type of SV is defined based on the location and orientation of the discordant reads that support the SV.

| Type of SV event | Definition |
| --- | --- |
| Deletion | Both breakpoints are within the 1 Mb window of the gene<br>Discordant reads at each breakpoint face inwards |
| Inversion | Both breakpoints are within the 1 Mb window of the gene<br>Discordant reads at each breakpoint face in same direction |
| Tandem duplication | Both breakpoints are within the 1 Mb window of the gene<br>Discordant reads at each breakpoint face outwards |
| Intergenic event | Only one breakpoint is within the 1 Mb window of the gene |

### Defining likely loss-of-function (LLoF) events

We classify SV events for a given gene as LLoF if they follow any of these three scenarios:

1. **SV interrupts the protein-coding part of a gene.** Parts of a gene being deleted or fused to another region of the genome such that the protein-coding portion of the gene is interrupted or truncated. For example, a scenario in which the intron between the second and third coding exons of a gene is fused with an intergenic region, such that the first two coding exons of the gene no longer connect to the latter coding exons, is considered an LLoF event. If, however, the SV results in an in-frame fusion with another gene, we do not consider it as an LLoF event.
2. **Out-of-frame fusion.** Occurs when two intronic/exonic regions are fused together such that their transcripts would be in opposite directions, or their transcripts’ reading frames would become out of frame with each other. This can occur between two genes, through an intergenic event, or within the same gene, through an intragenic inversion or tandem duplication. This likely results in translation of a non-functional protein, since a portion of the fused transcript would likely not correspond to a meaningful amino acid sequence. Furthermore, such out-of-frame fusions often code for a premature stop codon and induce nonsense-mediated mRNA decay (NMD),^47^ which degrades such out-of-frame fusion transcripts.
3. **Interruption of a gene’s transcription start site.** Occurs when an SV event spans the transcription start site of a gene and hence likely interrupts the transcription of the gene. For example, an intragenic SV event deleting part of the promoter and 5’UTR corresponding to a gene’s transcript may prevent the transcription initiation complex from properly being formed, thus leading to lower expression of the gene, likely disrupting its function.

From the above scenarios, we can categorize all possible types of SV events, either observed or modeled, that have at least one breakpoint within a gene’s neighborhood (i.e., within 1 Mb of the gene ends) as either likely causing the gene to lose its function (LLoF) or not (ULoF) (**Figures S1 and S2**).

**Figure S1:**
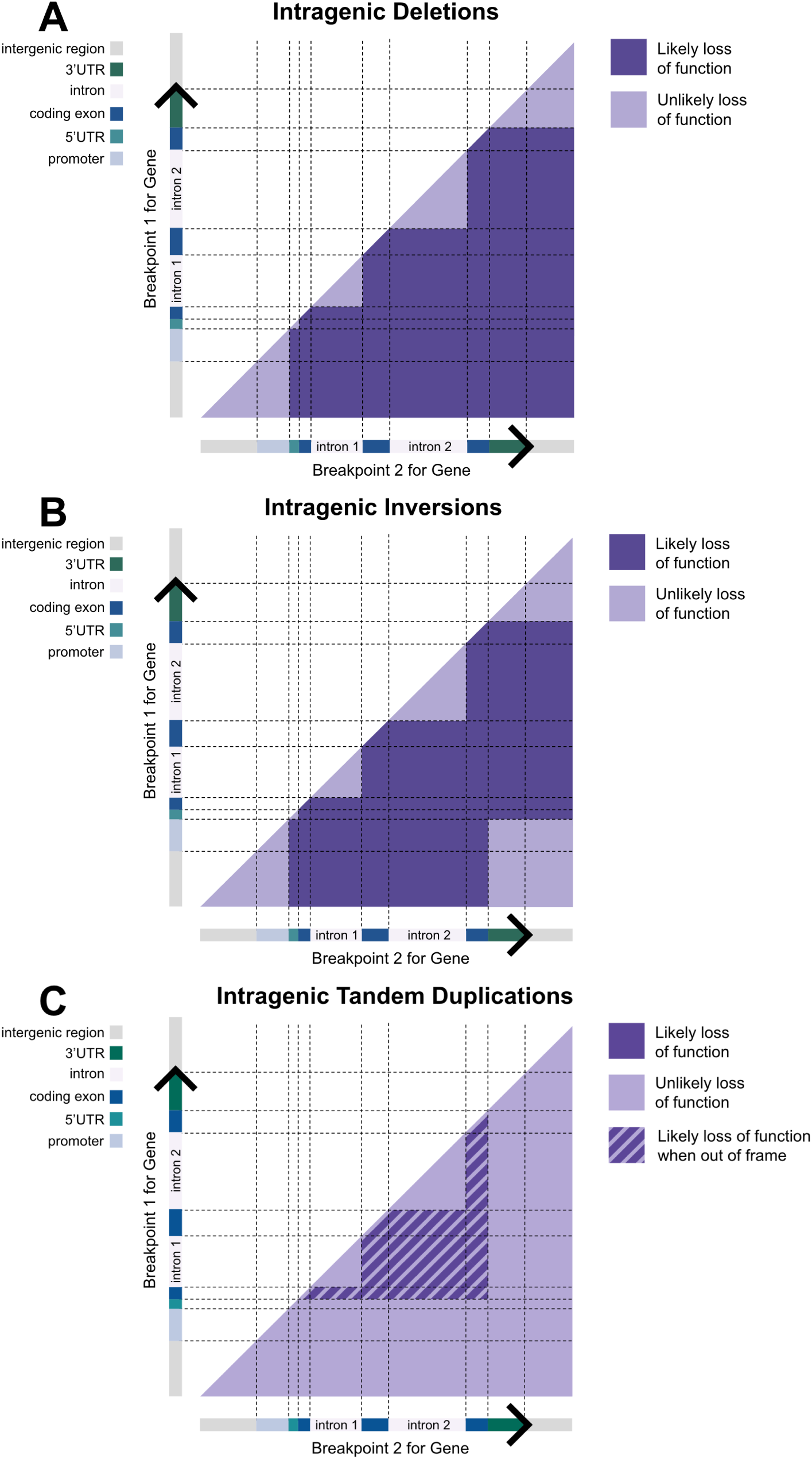
Visualization of LLoF conventions for intragenic events. For all panels, both axes represent different gene regions for the same gene. Each panel represents the LLoF conventions for intragenic SV events of different types: **A.** deletions, **B.** inversions, and **C.** tandem duplications. Note that all introns shown and numbered are between coding exons.

**Figure S2:**
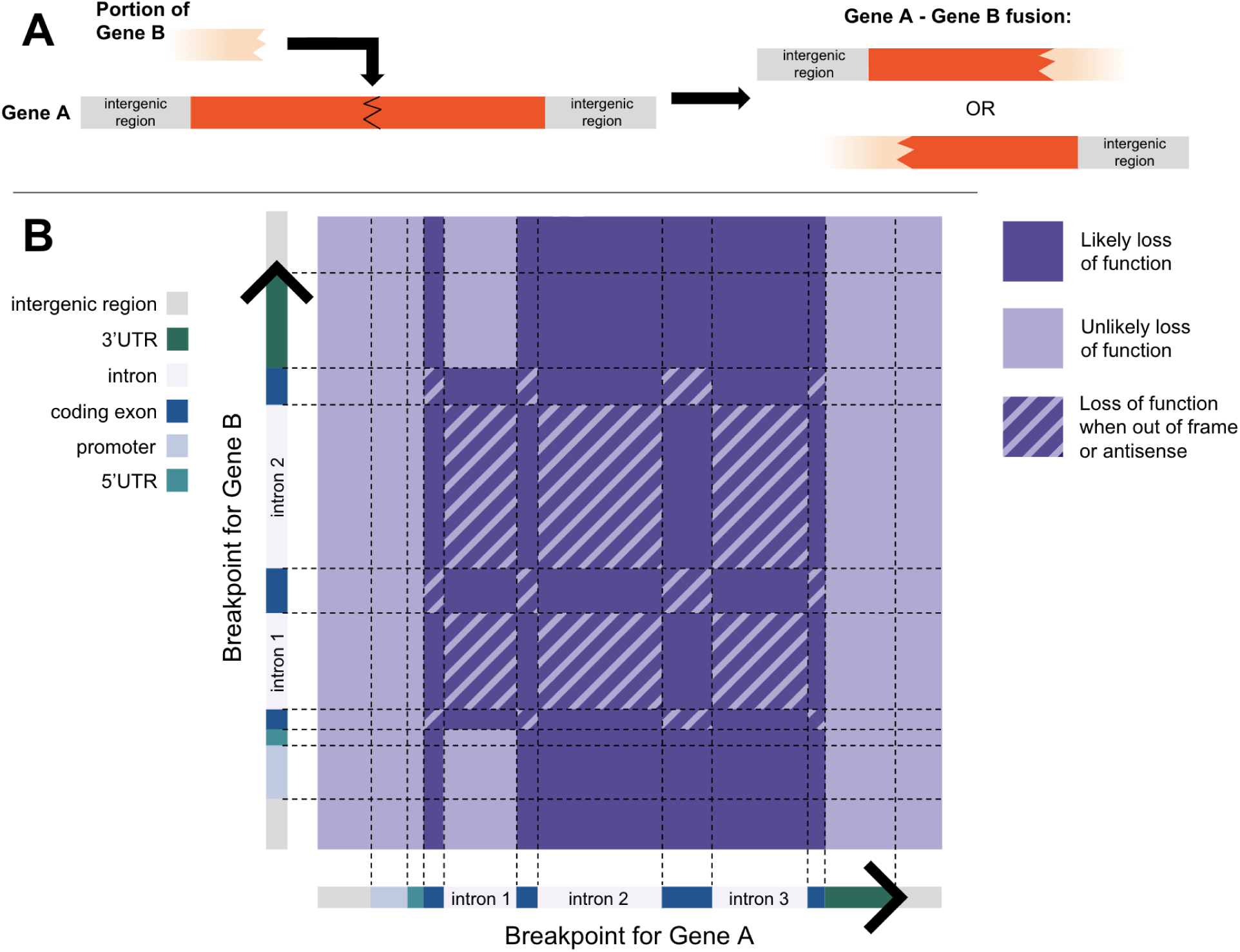
Visualization of LLoF conventions for intergenic SV events. **A.** An example of an intergenic SV event. A portion of a given gene (Gene B) has a breakpoint, and DNA from either side of it is incorporated into Gene A. **B.** All possible breakpoints from Genes A and B for an intergenic SV event, and an indication of whether the combination of locations of such breakpoints would result in LLoF for Gene A by our convention. Note that all introns shown and numbered are between coding exons. Moreover, the first intron between coding exons for the “collapsed” transcript for Gene A is treated differently since the first coding exon is often actually non-coding in overlapping transcripts, hence a disjunction of this first exon is considered a ULoF event.

### Annotating SV breakpoints

SVelfie connects SV breakpoints with genes in two ways: (i) it associates an SV breakpoint with all genes that are within a 1 Mb window (which can be an empty set), and (ii) it annotates each SV breakpoint with its closest gene (even if it is beyond the 1 Mb window). For annotations, we use a modified version of the annotator of dRanger^22^. The annotation algorithm is also used to infer the transcript domains in which breakpoints are observed in a given gene, from which we can determine whether certain SV events are LLoF or ULoF using the convention described above.

dRanger’s (and therefore SVelfie’s) annotator relies on the database of RefSeq^48^ hg18 transcripts (curated by the National Center for Biotechnology Information) used in the UCSC Genome Browser^49^. The transcripts were lifted over to hg38, and then processed to generate a table with attributes for each transcript, such as length, number of codons, exon boundaries, and exon frames. Relevant transcripts for certain genes were manually added as part of dRanger’s development, including *DUX4L1*, *HMGN2P46*, and *MALAT1*, as well as the mitochondrial genes *MT-CO2*, *MT-ATP8*, *MT-ATP6*, and *MT-ND4L*.

Using this table, the annotator finds the transcript which a given SV breakpoint would putatively cause the greatest functional effect in based on the type of region hit by the SV breakpoint and a functional severity ranking (**Table S2**). If there are multiple (often overlapping) transcripts that contain the SV breakpoint within the same type of region, the following rules apply: (i) For transcripts with breakpoints in promoter regions, the transcript with the breakpoint closest to one of the transcript edges is used; (ii) For transcripts with breakpoints in untranslated regions or introns, the transcript with the breakpoint closest to a coding exon is used; (iii) For transcripts with breakpoints in coding exons, the first listed transcript in the transcript table (derived from the original UCSC Genome Browser RefSeq transcript table) is used; and (iv) When a breakpoint does not overlap with any transcript, it is deemed to be in an intergenic region.

In order to annotate intergenic region breakpoints with a gene, the annotator searches further out until it identifies a transcript in the database. In practice, a window is iteratively increased around the SV breakpoint, and the database is queried for transcripts either before or after the breakpoint. If, during an iteration, multiple transcripts are found to be within the window, the transcript that is closest to the breakpoint is chosen.

**Table S2:** Order of severity of the functional effect based on genomic regions that are used in dRanger’s annotator.

| Order of functional effect | Genomic region |
| --- | --- |
| 1st | Coding exon |
| 2nd | Within coding intron |
| 3rd | 3' untranslated region (and enclosed introns) |
| 4th | 5' untranslated region (and enclosed introns) |
| 5th | Promoter (region within 3 kb of transcription start site) |
| 6th | Intergenic region |

For SVelfie, we modified the original dRanger annotator to prioritize genes from the Cancer Gene Census (v95; downloaded on December 13^th^, 2021) from the Catalogue of Somatic Mutations in Cancer (COSMIC)^44^ database that were indicated as being involved in translocations. The transcripts corresponding to any such genes were prioritized in their annotation over other genes, and among themselves were prioritized using the same prioritization described above, ensuring that known cancer driver genes related to driver SV events are prioritized by SVelfie’s annotator.

### Collapsing overlapping gene transcripts and annotating gene components

For each gene, we generate a “collapsed” transcript from the (often) multiple overlapping transcripts per gene, each of which has its own annotated genomic regions (e.g., 5’ UTR, coding exon 1, promoter, etc.). We did this by merging all the elements of all of the gene’s transcripts, and then annotating each region’s gene component by the transcript with the greatest (potential) functional effect (**Table S2**). Since different transcripts can start with different coding exons, or skip coding exons, it is easiest to define the regions where no coding exons are present but are still between coding exons found across any of the transcripts (i.e., the “within coding” intronic regions). We number the within coding intronic regions, starting with “1” for the intron closest to the transcription start site and increment the number as we advance along the transcript. The numbering is important for determining LLoF or ULoF based on our convention (**Figures S1 and S2**).

### SVelfie framework

A high-level description of the SVelfie model is shown in **Figure 1A**. SVelfie consists of four main steps:

1. **Filtering tested genes.** In order to limit the number of tested hypotheses and thus maximize power, we only analyze genes which are annotated to the breakpoints of at least two cases, and which have breakpoints associated to the gene across a minimum number of cases (here we used a threshold of 5).
2. **Calculating the expected fraction of LLoF events for each gene.** We represent the expected fraction of LLoF for a given gene as P(LLoF):

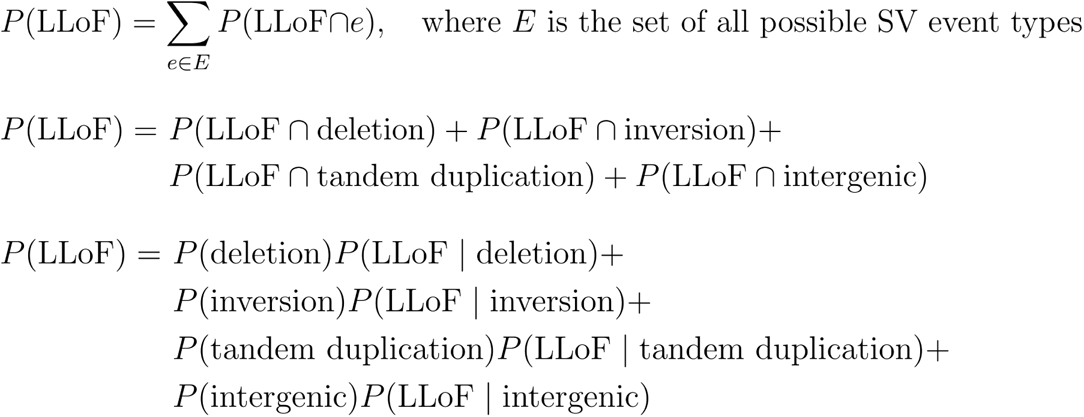 To calculate P(LLoF), we estimate each of the factors above, as follows:

- **P(deletion):** To estimate P(deletion), we use the proportion of deletion events across all SVs associated with genes annotated to the breakpoints of at least two cases. Note that the same SV event can be counted multiple times for different nearby genes.
- **P(inversion):** Perform the same process for P(deletion), but for inversions.
- **P(tandem duplication):** Perform the same process for P(deletion), but for tandem duplications.
- **P(intergenic):** Perform the same process for P(deletion), but for intergenic SV events.
- **P(LLoF | deletion):** To calculate P(LLoF | deletion), we assume the breakpoints are uniformly distributed within the 1 Mb window around the gene. The probability that a breakpoint occurs in a specific component of a transcript corresponds to the proportion of genomic territory that the particular component spans within the window of the gene (i.e., we assume the breakpoints follow a uniform distribution in this region). We also assume that the two deletion breakpoint locations are independent of one another, and hence the probability of the breakpoints landing in a certain combination of transcript components is simply the product of their probabilities. Hence, we have an estimate for the probabilities of all possible types of events given the event is intragenic. We can then match specific deletion events to whether or not they are indicated as LLoF in **Table S3** (as visualized in **Figure S1**), and sum up the associated probabilities of all such LLoF events to find P(LLoF | deletion).
- **P(LLoF | inversion):** Perform the same process for P(LLoF | deletion), but using the inversion LLoF conventions in **Table S3**.
- **P(LLoF | tandem duplication):** Perform the same process for P(LLoF | deletion), but using the tandem duplication LLoF conventions in **Table S3**.
- **P(LLoF | intergenic):** We use a similar process as for P(LLoF | deletion) — we also use the collapsed transcript for each gene and assume that both breakpoint locations are independent of one another. However, since only one breakpoint is inside of the 1 Mb window of the gene, we assume that that breakpoint has a uniform distribution within that window and that the other breakpoint has a uniform distribution across the entire genome. Thus, for the latter breakpoint, its probability of ending up in a certain transcript component is based on the proportion of that component across the entire genome. We estimate the genomic proportion of coding exonic regions by dividing the territory of the union of the coding exonic intervals of all transcripts (that are used by SVelfie) by the territory of the entire genome. To estimate the proportion of intronic regions between coding exons, we calculate the total genomic territory associated with introns that are between coding exons and divide by the genome length. With these estimates, we can calculate the probabilities for possible types of events given the event is intergenic by multiplying the proportions of corresponding gene components for the local breakpoint and the genome-wide non-local breakpoint. We can then calculate P(LLoF | intergenic) using **Table S4** (as visualized in **Figure S2**) to sum up the probabilities for all corresponding LLoF events.
3. **Calculating significance.** We use the conventions defined in **Tables S3** and **S4** for intragenic and intergenic SV events, respectively, to denote which observed events in the vicinity of each gene across the cohort are LLoF or ULoF. For each gene, we then use a binomial test with the observed number of LLoF events, the total number of SVs associated with the gene, and P(LLoF) (i.e., the expected fraction of LLoF events for each gene) to compute significance (i.e., p-value) of the gene.
4. **Correcting for multiple hypothesis testing.** We apply the Benjamini-Hochberg false discovery rate (FDR) correction procedure^50^ to correct for multiple hypothesis testing and calculate a q-value for each gene. Genes with q≤0.1 are considered significant, and genes with 0.1<q≤0.25 are considered near significant. These genes are SVelfie’s candidate drivers.

**Table S3:** Conventions used by SVelfie for LLoF versus ULoF for all possible intragenic SV events by genomic regions of breakpoints.

| Placement of breakpoints | Deletion cases | Tandem duplication cases | Inversions cases |
| --- | --- | --- | --- |
| before gene-before gene | ULoF | ULoF | ULoF |
| before gene-promoter | ULoF | ULoF | ULoF |
| before gene-5'UTR | LLoF | ULoF | LLoF |
| before gene-coding exon | LLoF | ULoF | LLoF |
| before gene-within coding intron | LLoF | ULoF | LLoF |
| before gene-3'UTR | LLoF | ULoF | ULoF |
| before gene-after gene | LLoF | ULoF | ULoF |
| promoter-promoter | ULoF | ULoF | ULoF |
| promoter-5'UTR | LLoF | ULoF | LLoF |
| promoter-coding exon | LLoF | ULoF | LLoF |
| promoter-within coding intron | LLoF | ULoF | LLoF |
| promoter-3'UTR | LLoF | ULoF | ULoF |
| promoter-after gene | LLoF | ULoF | ULoF |
| 5'UTR-5'UTR | ULoF | ULoF | ULoF |
| 5'UTR-coding exon | LLoF | ULoF | LLoF |
| 5'UTR-within coding intron | LLoF | ULoF | LLoF |
| 5'UTR-3'UTR | LLoF | ULoF | LLoF |
| 5'UTR-after gene | LLoF | ULoF | LLoF |
| coding exon-coding exon (in-frame) | LLoF | ULoF | LLoF |
| coding exon-coding exon (out-of-frame) | LLoF | LLoF | LLoF |
| coding exon-within codingintron (in-frame) | LLoF | ULoF | LLoF |
| coding exon-within coding intron (out-of-frame) | LLoF | LLoF | LLoF |
| coding exon-3'UTR | LLoF | ULoF | LLoF |
| coding exon-after gene | LLoF | ULoF | LLoF |
| within coding intron-within coding intron (in-frame) | LLoF | ULoF | LLoF |
| within coding intron-within coding intron (out-of-frame) | LLoF | LLoF | LLoF |
| within coding intron-within coding intron (within same intron) | ULoF | ULoF | ULoF |
| within coding intron-3'UTR | LLoF | ULoF | LLoF |
| within coding intron-after gene | LLoF | ULoF | LLoF |
| 3'UTR-3'UTR | ULoF | ULoF | ULoF |
| 3'UTR-after gene | ULoF | ULoF | ULoF |
| after gene-after gene | ULoF | ULoF | ULoF |

**Table S4:** Conventions used by SVelfie for LLoF versus ULoF for a given gene Gene A for all possible intergenic SV events by genomic regions of breakpoints (where Gene B is some other distally located gene).

| Breakpoint placement relative to Gene A | Breakpoint placement relative to Gene B | In-frame or out-of-frame/antisense | Likely LoF (LLoF) or unlikely LoF (ULoF) for Gene A |
| --- | --- | --- | --- |
| intergenic region | intergenic region | N/A | ULoF |
| intergenic region | promoter | N/A | ULoF |
| intergenic region | 5'UTR | N/A | ULoF |
| intergenic region | coding exon | N/A | ULoF |
| intergenic region | within coding intron | N/A | ULoF |
| intergenic region | 3'UTR | N/A | ULoF |
| promoter | intergenic region | N/A | ULoF |
| promoter | promoter | N/A | ULoF |
| promoter | 5'UTR | N/A | ULoF |
| promoter | coding exon | N/A | ULoF |
| promoter | within coding intron | N/A | ULoF |
| promoter | 3'UTR | N/A | ULoF |
| 5'UTR | intergenic region | N/A | ULoF |
| 5'UTR | promoter | N/A | ULoF |
| 5'UTR | 5'UTR | N/A | ULoF |
| 5'UTR | coding exon | N/A | ULoF |
| 5'UTR | within coding intron | N/A | ULoF |
| 5'UTR | 3'UTR | N/A | ULoF |
| coding exon | intergenic region | N/A | LLoF |
| coding exon | promoter | N/A | LLoF |
| coding exon | 5'UTR | N/A | LLoF |
| coding exon | coding exon | in-frame | ULoF |
| coding exon | coding exon | out-of-frame or antisense | LLoF |
| coding exon | within coding intron | N/A | LLoF |
| coding exon | 3'UTR | N/A | LLoF |
| within coding intron 1 | intergenic region | N/A | ULoF |
| within coding intron 1 | promoter | N/A | ULoF |
| within coding intron 1 | 5'UTR | N/A | ULoF |
| within coding intron 1 | coding exon | N/A | LLoF |
| within coding intron 1 | within coding intron | in-frame | ULoF |
| within coding intron 1 | within coding intron | out-of-frame or antisense | LLoF |
| within coding intron 1 | 3'UTR | N/A | ULoF |
| within coding intron 2+ | intergenic region | N/A | LLoF |
| within coding intron 2+ | promoter | N/A | LLoF |
| within coding intron 2+ | 5'UTR | N/A | LLoF |
| within coding intron 2+ | coding exon | N/A | LLoF |
| within coding intron 2+ | within coding intron | in-frame | ULoF |
| within coding intron 2+ | within coding intron | out-of-frame or antisense | LLoF |
| within coding intron 2+ | 3'UTR | N/A | LLoF |
| 3'UTR | intergenic region | N/A | ULoF |
| 3'UTR | promoter | N/A | ULoF |
| 3'UTR | 5'UTR | N/A | ULoF |
| 3'UTR | coding exon | N/A | ULoF |
| 3'UTR | within coding intron | N/A | ULoF |
| 3'UTR | 3'UTR | N/A | ULoF |

### Input data

The only user input necessary to run SVelfie is a tab-separated values file (.tsv) of the SVs detected across a cohort (with each row containing the information for a single SV). These are the required 10 columns and their headers (the order of the columns does not matter):

- “individual”: An identifier for a particular tumor case.
- “chr1”: The integer chromosome number of the upstream breakpoint (breakpoint 1) for a particular SV. Chromosome X is represented as “23”, and chromosome Y is represented as “24”.
- “str1”: The strand (or orientation) of the discordant reads supporting breakpoint 1 for a particular SV. “0” signifies they are forward-facing, meaning that genomic content just upstream of the breakpoint (according to the reference genome) is involved in the given SV. “1” signifies they are in the opposite strand, reverse-facing, meaning that genomic content just downstream of the breakpoint (according to the reference genome) is involved in the given SV.
- “pos1”: The one-based (first base of chromosome is numbered 1) position of breakpoint 1 within a chromosome for the more upstream breakpoint for a particular SV.
- “chr2”: The integer chromosome number of the downstream breakpoint (breakpoint 2) for a particular SV, as for breakpoint 1.
- “str2”: The strand information for breakpoint 2, as for breakpoint 1.
- “pos2”: The one-based position of breakpoint 2, as for breakpoint 1.
- “span”: The difference between pos1 and pos2 for a particular intrachromosomal SV; otherwise, the value is blank.
- “gene1”: The annotated gene for breakpoint1 using SVelfie’s annotator.
- “gene2”: The annotated gene for breakpoint2 using SVelfie’s annotator.

### RNA-seq depletion statistical test

For each candidate driver gene, to test for lower RNA-seq expression in the cases with SVs, we used a one-sided Wilcoxon rank sum test between the TPM-corrected RNA-seq values of patient cases that had at least one SV breakpoint within a 1 Mb vicinity of the gene and those that did not (using patient cases where such RNA-seq data is available). We corrected for multiple hypothesis testing (across all candidate drivers) using the Benjamini-Hochberg procedure to calculate q-values.

### Confidence intervals for observed LLoF fractions

Confidence intervals for observed LLoF fractions in **Figure 2A** were calculated via a beta distribution, reflecting the posterior distribution of the fraction given the observed counts and a uniform prior.

### Q–Q plot p-value correction

For displaying the Q–Q plot, a correction was applied for the displayed p-values in **Figure 2B** to make their distribution continuous and uniform under the null hypothesis; we did this by uniformly randomly sampling between the original p-value and the next more significant p-value that corresponds to the next discrete value in the binomial test. The bounds of the interval for sampling for each gene are represented by vertical lines.

### Dataset

SVelfie was run on the cohort of patient cases with “lymph nodes” as their afflicted organ system in PCAWG’s clinical specimen histology table. There were 107 patient cases total, 106 of which had at least one SV.

### Code

SVelfie is publicly available for use at https://github.com/getzlab/SVelfie. Directions for running SVelfie can be found in the file run/directions_for_running_model.txt within the repository.

## Acknowledgments

This work was partially funded by a collaboration with Inocras, Inc. (to G.G. and E.R.), as well as a Broad-internal SPARC award (to G.G. and E.R.). G.G. was partially funded by the Paul C. Zamecnik Chair in Oncology at the Mass General Cancer Center. We thank Getz Lab and Rheinbay Lab members for their feedback and ideas for SVelfie, and Rheinbay Lab members for their feedback on the manuscript figures (especially Philipp Hahnel and Preshita Dave).

